# Discovering genetic biomarkers for targeted cancer therapeutics with eXplainable AI

**DOI:** 10.1101/2023.07.24.550346

**Authors:** Debaditya Chakraborty, Elizabeth Gutierrez-Chakraborty, Cristian Rodriguez-Aguayo, Hakan Başağaoğlu, Gabriel Lopez-Berestein, Paola Amero

## Abstract

Explainable Artificial Intelligence (XAI) enables a holistic understanding of the complex and nonlinear relationships between genes and prognostic outcomes of cancer patients. In this study, we focus on a distinct aspect of XAI – to generate accurate and biologically relevant hypotheses and provide a shorter and more creative path to advance medical research. We present an XAI-driven approach to discover otherwise unknown genetic biomarkers as potential therapeutic targets in high-grade serous ovarian cancer, evidenced by the discovery of IL27RA, which leads to reduced peritoneal metastases when knocked down in tumor-carrying mice given IL27-siRNA-DOPC nanoparticles.

**Summary:** Explainable Artificial Intelligence is amenable to generating biologically relevant testable hypotheses despite their limitations due to explanations originating from post hoc realizations.

## Main

High-grade serous ovarian cancer (HGSC) is the most frequent and leading cause of death from gynecologic cancers (*1*) that accounts for 70–80% of ovarian cancer deaths, and overall survival has not improved significantly for several decades (*2*). Patients diagnosed with advanced HGSC have a 5-year survival rate of 41% (*3*), analysis of which could provide insights into tumor biology and therapeutic approaches (*4*). Taking notes from previous studies on HGSC, we present an integrated eXplainable Artificial Intelligence (XAI) and probabilistic approach on HGSC data (*n=407, where n is the sample size*) to identify inherent relationships between the genetic features and the ≥5-year survival probability. The objective of this article is to uncover the most critical biomarkers from a pool of 655 potential targets using our unique data-driven approach.

Recent studies indicate that AI models are often referred to as ‘black boxes’because their decisionmaking process lacks transparency (*5*). The consensus is that the lack of inherent explainability is problematic as this produces biases, creates difficulties in detecting false positives and negatives and hides potential insights that may be derived from AI (*6*). In this study, we provide evidence highlighting the potential of XAI in enhancing biological explainability by uncovering novel insights from the underlying data. Our XAI approach predicts patient outcomes and survival duration based on genetic signatures (predictive AI aspect of the models) and discover & visualize critical biomarkers (biological explainability aspect of the models) in HGSC. To ensure the viability of explanations generated by our XAI, we subsequently validated the most prominent HGSC promoting biomarker identified by XAI using in-vivo murine tumor models (schematic shown in Supplementary Text 1). The XAI approach outlined in this study is not only intended to generate high predictive accuracy but also infer the cause-effect relations behind the predictions and identify counterfactuals that are useful for optimizing interventional therapies and assess the resultant improvements in patients.

Let us briefly discuss a hypothetical situation to further clarify the benefit of an explainable AI approach in biomedical domains: Imagine having to choose between a surgeon who could explain anything about the surgery but had a 15% chance of causing death during the surgery or a robotic arm that could perform the surgery with only a 2% chance of failure but will not explain anything about the surgery – most people choose the surgeon with 85% success rate (*7*). Integrating high-performance AI models with post hoc explainability methods can bring about the best of both worlds – the knowledge & wisdom of the surgeon along with the precision of the robotic arm. In this paper, we show that one does not necessarily have to sacrifice between predictive performance (e.g., accuracy, precision, and recall) and explainability (from a biological point of view) that can generate fresh insights.

We report that our models predicted the ≥5-year survival probability from genetic features of patients (*n=407*) with 97.52% accuracy, 100% precision, and 94.74% recall on the testing data that comprised of the 25% of the total samples that were hidden from the models during the training phase. Insights derived through XAI prioritized the biomarkers that are of utmost importance in determining prognosis for patients with HGSC, which we refer to as ‘global biological explanations’(shown in Figure 1 A). The biomarkers are arranged in terms of their relative importance (most important at the top), and the variations in concentration of the biomarkers from high (red) to low (blue) together with the corresponding Shapley values – on the x-axis – are used to determine whether a particular biomarker is associated with poor (positive values on the x-axis) or good (negative values on the x-axis) prognosis in HGSC (Figure 1 A). For example, we discovered that TAF10 is associated with good prognosis since higher concentrations (red) of this biomarker correspond to negative Shapley values on the x-axis – hence it can be classified as a cancer dampener for HGSC patients. In contrast, IL27RA is associated with poor prognosis since higher concentrations (red) of this biomarker correspond to positive Shapley values on the x-axis – hence it can be classified as a cancer promoter for HGSC patients. Furthermore, XAI unveiled the inflection points of each influential gene (local biological explanations) above or below which the prognosis would either improve or deteriorate (Figure 1 B). These inflection points serve as guidance for genomic editing that could lead to the development of new gene therapeutics and enhanced prognosis of patients.

**Figure 1:**
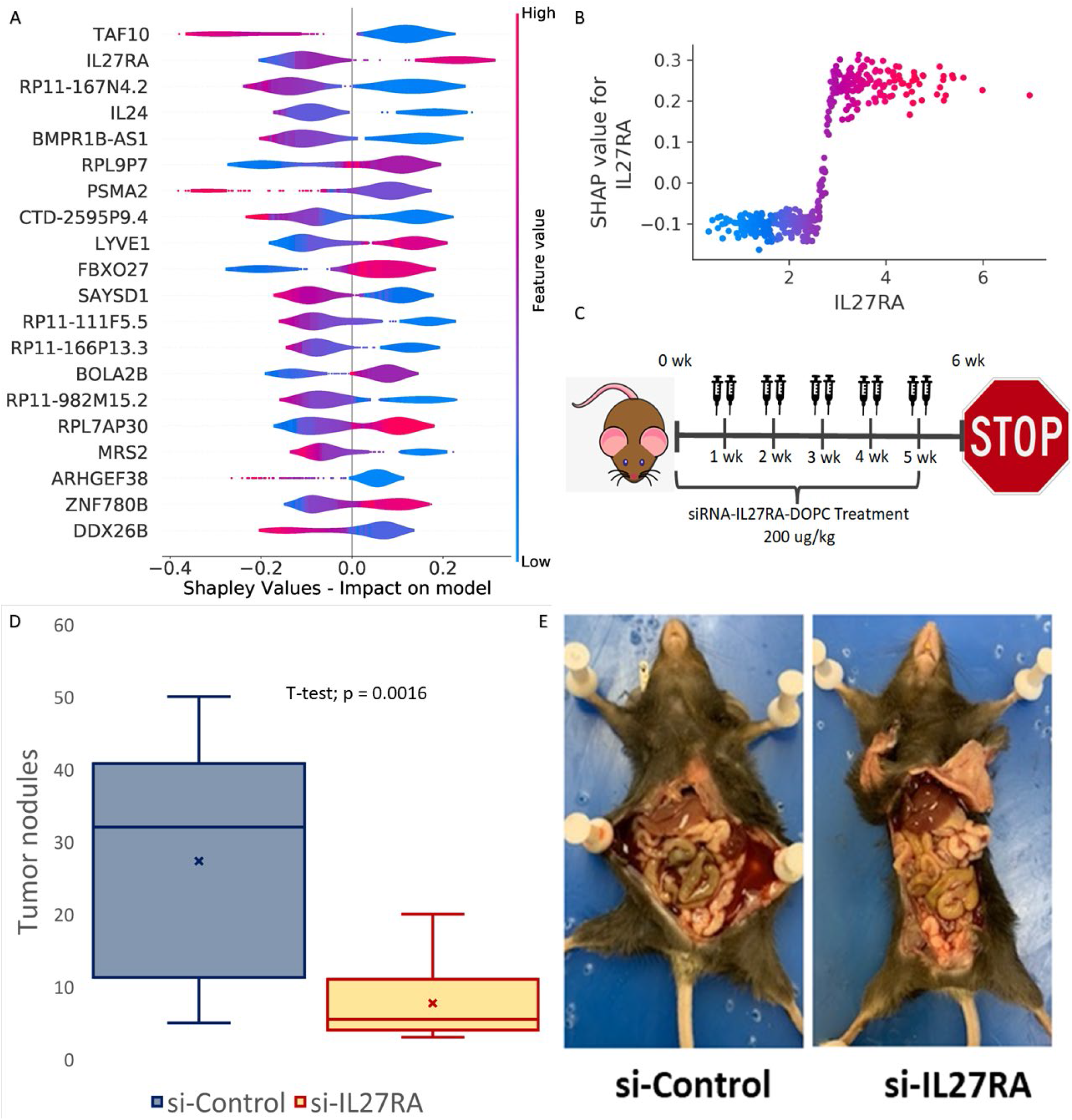
(A) Global biological explanations produced by XAI, (B) local biological explanations for the most prominent cancer promoting biomarker (IL27RA) gene produced by XAI, (C) schedule of treatment of si-IL27RA in vivo model, (D) statistical comparison of metastasis (represented by tumor nodules) in the control group and mice given si-IL27RA treatment, and (E) representative pictures of ID8 tumor bearing Cbearing C57BL/6 mice.

Based on insights generated by XAI, we discovered that IL27RA (pathway analysis in Supplementary Text 2) is the most prominent cancer promoting biomarker associated with lower survival probabilities in HGSC patients. Based on these XAI results, we hypothesize that editing the concentration of IL27RA below 2.2 TPM would lead to improved prognosis. To validate this XAI-driven hypothesis, we conducted a study in an orthotopic model of ovarian cancer in C57BL/6 mice intraperitoneally (IP) inoculated with one million ID8 ovarian cancer cells. This model shows extensive intraperitoneal dissemination consistent with the human version of the disease. Two groups of 10 mice, each were injected IP with the tumor cells, group one was injected IV with DOPC nanoparticles (NP) loaded with scrambled siRNA and group 2 was injected with DOPC-NP loaded with IL27-siRNA (IV) Figure 1 C-E. The two treatments were administered IV twice a week for 5 weeks, at which time the mice were sacrificed and peritoneal metastasis were counted. The results, in Figure 1 D and E, show a significant reduction in peritoneal metastasis in mice injected with the IL27-siRNA-DOPC NP. The findings from this study suggest that silencing IL27RA will result in enhanced prognosis of HGSC patients.

The biology of IL27 is complex and context dependent – it may either dampen or promote different types of inflammation or cancer (*8*). Recently, an experimental study revealed that IL27 receptor signaling promotes hepatocellular carcinoma, and high expression correlated with poor prognosis for patients due to increased proliferative capacity of tumors and inflammation (*9*) - thus further validating that the explanations obtained from XAI are, in fact, legitimate.

A challenge is that lab experiments, to corroborate findings from XAI, are time and resource-consuming. Thus, we describe a probabilistic approach to evaluate insights unraveled from the XAI models relating to the underlying relationships between the features and targets (Materials and Methods). The probabilistic approach shows - from a data-driven perspective - whether the associations highlighted by XAI are valid from an interventional therapeutics standpoint (Figure 2). Based on causality estimated with the probabilistic approach, we found that modulating the top five biomarkers combinatorically (Figure 1 A and Figure 2 A-F) can improve the chance of survival in HGSC patients (beyond five years) by 100%. Interestingly, TATA-box binding protein associated factor 10 (TAF10), identified as the leading HGSC suppressor by our XAI (Figure 1 A and Figure 2 A & F), can probabilistically cause 41.1% improvement individually. The literature further indicates that TAF10 may have significant implications, such as enhancing tumor-suppressing p53 activity (*10*–*15*) and regulating MYC transcriptional activity (*16, 17*). However, further studies are required to advance the mechanistic link (if any) between TAF10 and tumor suppression. Kaplan-Meier and box-plot analysis of TAF10 and IL27RA also corroborate our XAI-based findings, through which we observe an increase in survival chances corresponding to higher expressions of TAF10, while higher expression of IL27RA is associated with reduced chances of survival (Supplementary Text 3). Based on the evidence from this study, XAI approaches are amenable to generating biologically relevant testable hypotheses despite their limitations due to explanations originating from post hoc realizations (*18*). Probabilistic causality approaches, as described in this study, can further reinforce findings from XAI and provide a shorter path to advance precision oncology while saving time and money.

**Figure 2:**
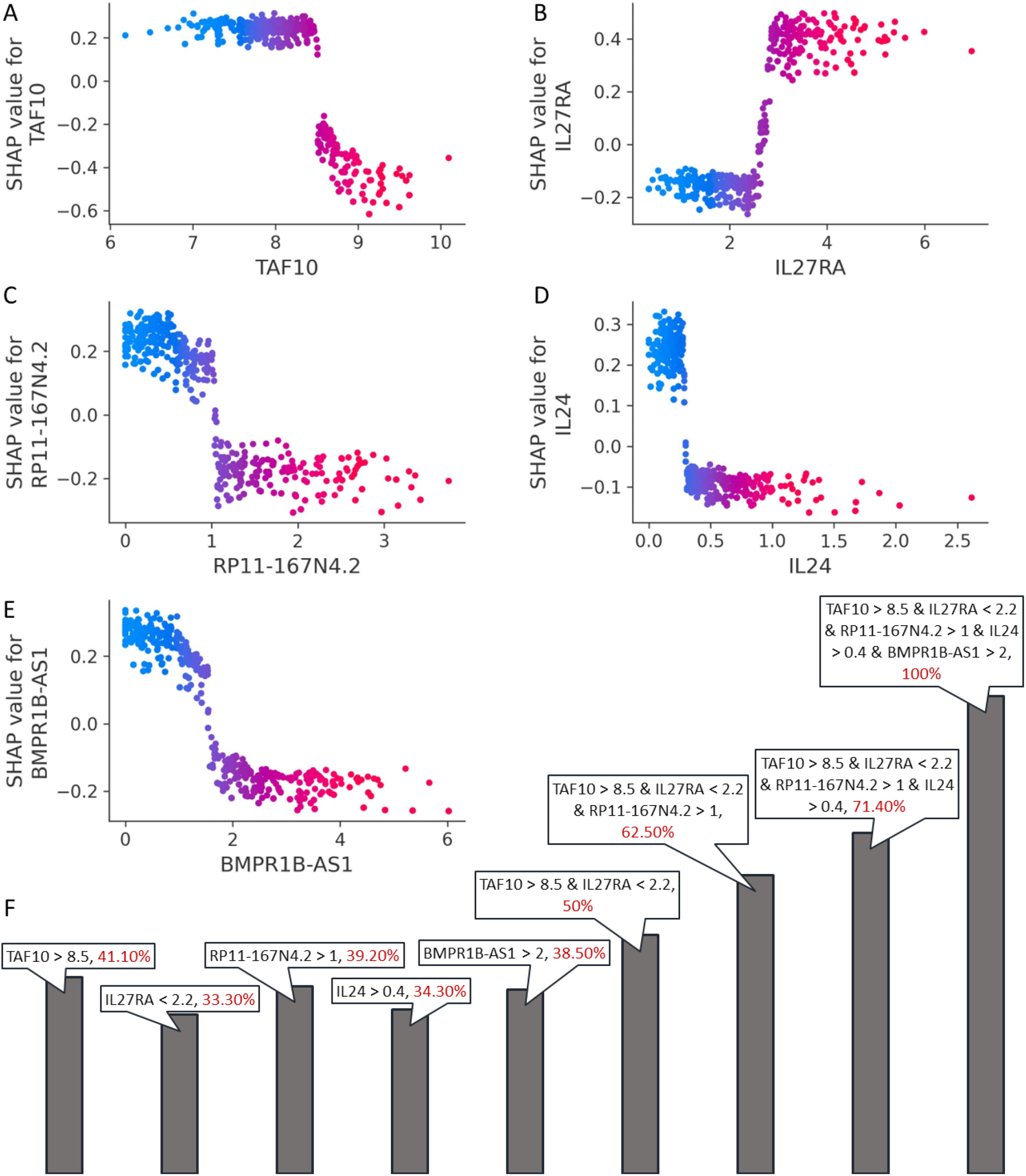
(A-E) Local biological explanations for the top five critical biomarkers identified by XAI. (F) Probability of improvements in therapeutic outcome when the biomarkers are either individually (or combinatorically) targeted in the tumor microenvironment.

In this study, we exclusively focused on designing our integrated XAI and probabilistic approach and experimentally testing the explanations - generated by XAI - associated with IL27RA using in-vivo murine tumor models. Future work on the application of XAI to advance cancer research could shape up in three ways: (I) novel implementations of our approach to other cancer types and algorithmic & data specific improvements; (II) experimentally testing the viability of explanations through XAI for other critical biomarkers besides IL27RA; (III) investigating novel ways of implementing the knowledge gained from XAI and integrating into clinical therapeutics. Leveraging the integrated XAI and probabilistic approach, described in this study, to identify genetic biomarkers relevant to patients and nanoparticles to provide targeted anti-cancer precision therapy is a cost- and time-efficient technique that offers an innovative and effective solution for the treatment of different types of diseases.

## Data Availability

The datasets and code for this study will be published in Code Ocean once accepted for publication. Links will be shared at that time.

## Acknowledgements

We thank Cristina Ivan for helping with data acquisition while at MDACC. This study was supported in part by grants from the National Institutes of Health/National Cancer Institute (5U01CA213759-02, P30CA016672), and the American Cancer Society, National Science Foundation (CHE-1411859), and an endowment grant from the John P. Gaines Foundation. Dr. Cristian Rodriguez-Aguayo and Dr. Paola Amero were supported by the Brain SPORE Career Enhancement Program and NCI grant P50CA127001, as well as by the NIH through the Ovarian SPORE Career Enhancement Program and NCI grant P50CA217685.

## List of Supplementary Materials

Materials and Methods

Supplementary Text (1-3)

Figs. S1 to S3

References (1–17) in Supplementary Materials.

## Author Contribution

Conceived and designed the experiments: D.C., H.B., and G.L.B.; Performed the experiments: D.C., P.A., and CRA; Analyzed the data: All authors; Contributed materials/analysis tools: All authors; Wrote the paper: D.C. and E.G.C. primarily and all authors reviewed, revised, and commented on the manuscript.

## Materials and Methods

The data used for this pilot study primarily originates from TCGA Ovarian cancer cohort (*1*), immune cell fraction for different cell types as estimated by EPIC (*2*) along with expression for certain pathway of interest. The data that we worked with composed of clinical information from cbioPortal (*3*), PanCancerAtlas (*4*), and RNASeq expressions for the corresponding cohort quantified as TPM from UCSC XENA project (*5, 6*).

The XAI model design, motivated from XGBoost (*7*) and SHAP (*8*) frameworks, in this study can be represented by the equation: Ŷ = *f*(*X*^*1*^, *X*^*2*^, …, *X*^*n*^); where Ŷ represents the predicted outcomes, *f* is the functional representation of the relationship between *X*(i.e., the independent features – biomarkers) and Ŷ (i.e., target – the true outcome). The function *f*is learnt by minimizing a regularized objective function that consists of two parts: the loss function (*L*(*y*_*i*_,ŷ_*i*_)) that measures the difference between the predicted outcome ŷ_*i*_ using the weighted ensemble of regression trees and the true outcome y_*i*_, and the regularization term (Ω(*w*)) that penalizes complex models for a given set of *w* (vector of weights assigned to the individual regression trees). The regression trees are iteratively added to the ensemble by computing the negative gradient of the objective function with respect to the current ensemble predictions 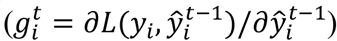 where 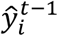 is the predicted outcome of the ensemble in iteration *t* − *1*. The negative gradient represents the direction of steepest descent for the objective function, and it indicates how much the current ensemble output needs to be corrected to minimize the objective function. The model then fits a new tree *h*_*t*_(*x*) to the negative gradient values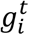, by minimizing an objective function 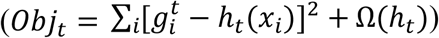 is the predicted outcome of the *t*^*th*^ tree for the input *x*_*i*_ and Ω(*h*_*t*_) is the regularization term that penalizes complex trees. The optimal weights for the new tree *h*_*t*_*(x*) are computed using 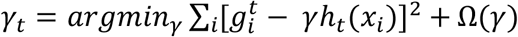); where *γ*_*t*_ is the optimal weight for the new tree *h*_*t*_*(x*) and Ω(γ) is the regularization term that penalizes large weights. The model then updates the predicted outcome by adding the new tree with weight 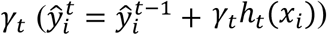. This process is repeated until the objective function converges. The final predicted outcomes of the model are the weighted sum of the predicted outcomes from all the trees in the ensemble. In this study, we found that 5000 regression trees with a maximum depth of 5, learning rate of 0.01, and 60% subsampling generates optimal results based on a grid-search crossvalidation approach. Feature attribution is computed using Shapley values by introducing each feature, one by one, into a conditional expectation function of the model’s predicted outcome (*f*_*x*_(*S*) = *E*[*f*(*X*)|*do*(*X*_*s*_ = *x*_*s*_)), and attributing the change produced at each step to the feature that was introduced and then averaging this process over all possible feature orderings; where *S* is the set of features conditioned on, *X* is a random variable representing the model’s *M* input features, *x* is the model’s input vector for the current prediction.

A grand challenge is that XAI approaches in medical domains are widely speculated because the insights are purely associative (*9, 10*). Therefore, we propose a probabilistic causal inference theory to evaluate the impacts of insights unraveled from the XAI models relating the underlying relationships between the features and the targets. Through this approach, the likelihood of improvements in the targets is determined based on increases/decreases in values of the features in reference to feature attribution and Shapley interaction index from game theory – which are analyzed to mathematically assess the effectiveness of specific therapeutic interventions. We formulated the probabilistic models to infer causality from the associative links revealed with XAI by using 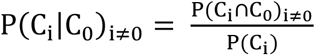 for individual cancer promoters/suppressors and 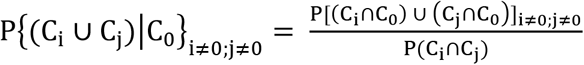 for combinatorial sets of cancer promoters/suppressors; where *C0* is the number of patients in the cohort that survived (≥5 years), *C*_*i*_ and *C*_*j*_ are the number of patients in the cohort that satisfy the genetic conditions (*i* & *j*) identified by XAI, and *P*(*C*_*i*_ *∩ C*_*j*_) is the probability that conditions *i* & *j* are both satisfied.

### Supplementary Text 1

**Figure S1:**
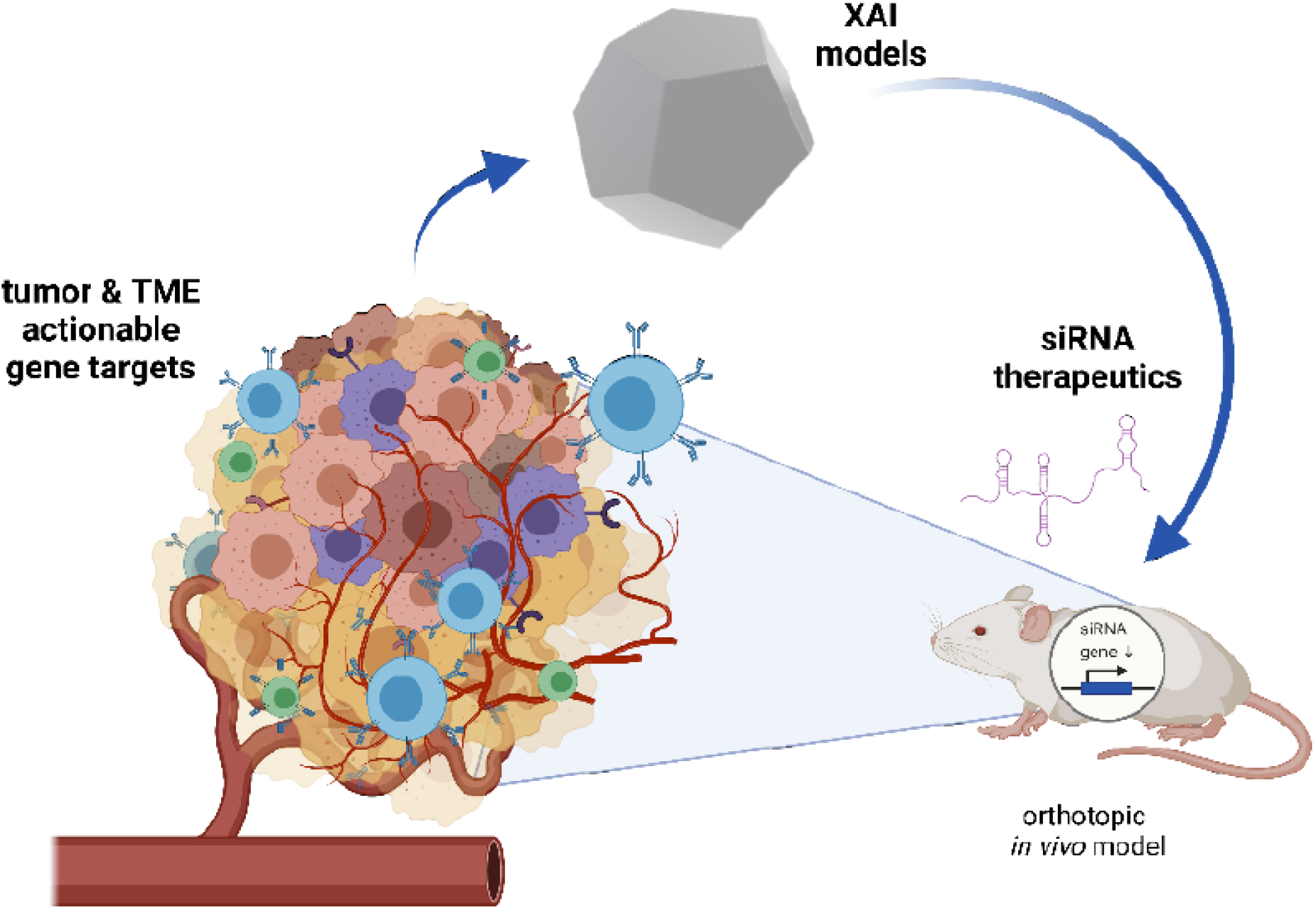
Schematic of an integrated XAI approach with orthotopic in vivo model to identify otherwise unknown critical genes in the tumor and tumor microenvironment (TME) for therapeutic targets

### Supplementary Text 2

IL-27RA is a receptor for Interleukin-27 (IL-27) (*11*), a heterodimeric cytokine made up of Epstein Barr virus induced gene 3 (EBI_3_) and p28 (*12*). The IL-27RA can be found on CD4+ cells, which then differentiates into TH1 cells (*13*). Through this pathway the cells secrete IFN-gamma (a proinflammatory cytokine). It is not until further in the pathway (Figure S2) that IL-10 (an antiinflammatory cytokine) is secreted. Thus, increased IL-27RA expressions could lead to chronic inflammation causing an increase in metastasis and consequently poor prognosis in HGSC patients. Inflammation has been considered a hallmark trait of cancer (*14*–*16*). Chronic inflammation introduces an environment where cancer can grow and metastasize. Inflammation causes an increase in blood flow and nutrients to be brought to the affected area. Chronic inflammation is an important high-risk factor for HGSC as well as epithelial ovarian cancer (*17*). IL-27RA has been mostly overlooked in favor of IL-27, but due to advances in XAI techniques, IL-27RA has been found to be a critical biomarker for the potential treatment of HGSC.

**Figure S2:**
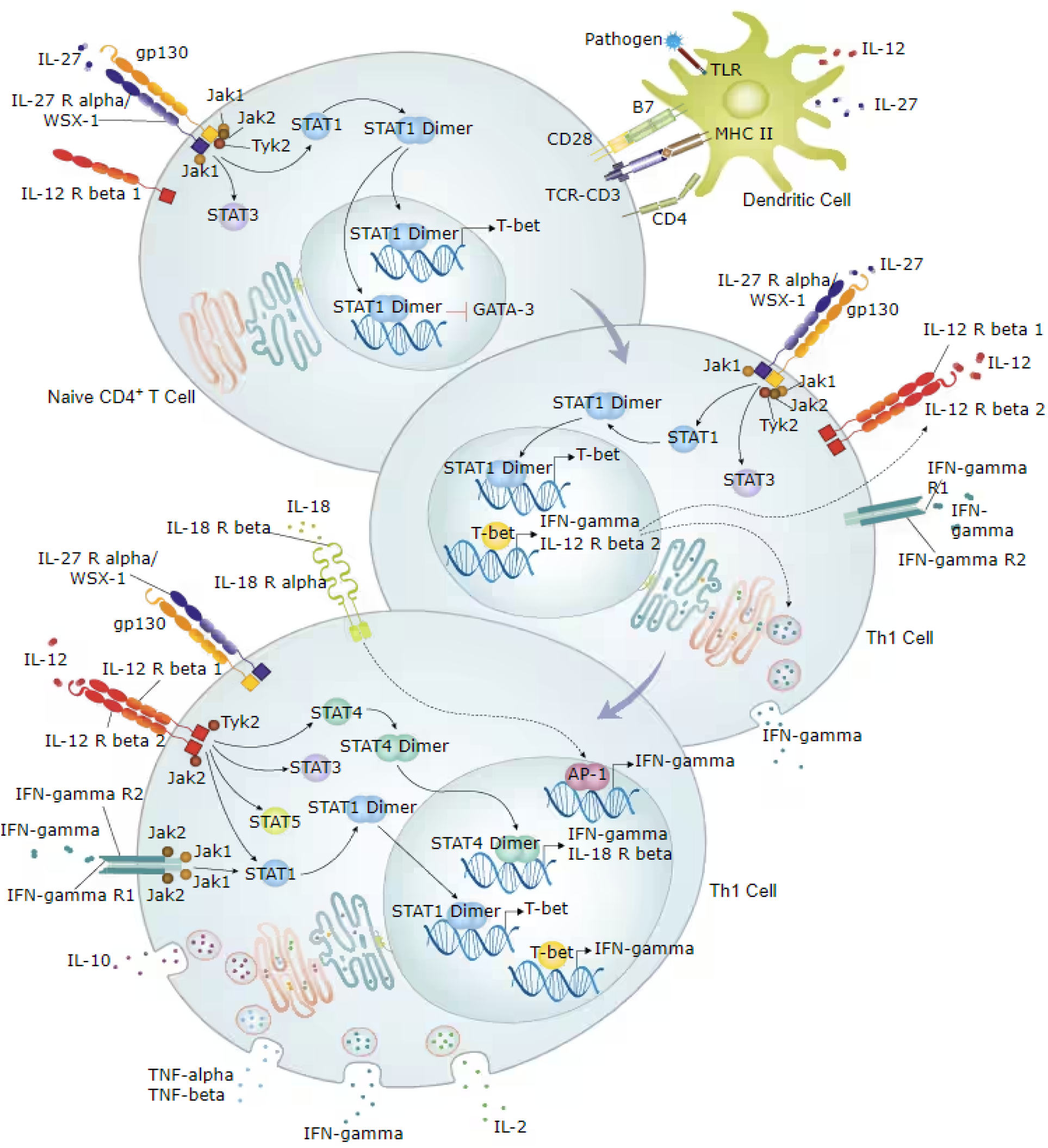
TH1 Pathway representation to understand how IL27RA relates to poor prognosis in HGSC. Source: https://www.rndsystems.com/pathways/th1-differentiation-pathway

### Supplementary Text 3

Shapley, Kaplan-Meier, and box-plot analyses of TAF10 and IL27RA demonstrate that an increase in survival chances corresponding to higher expressions of TAF10, while higher expression of IL27A is associated with a reduction in chances of survival of HGSC patients (Figure S3).

**Figure S3:**
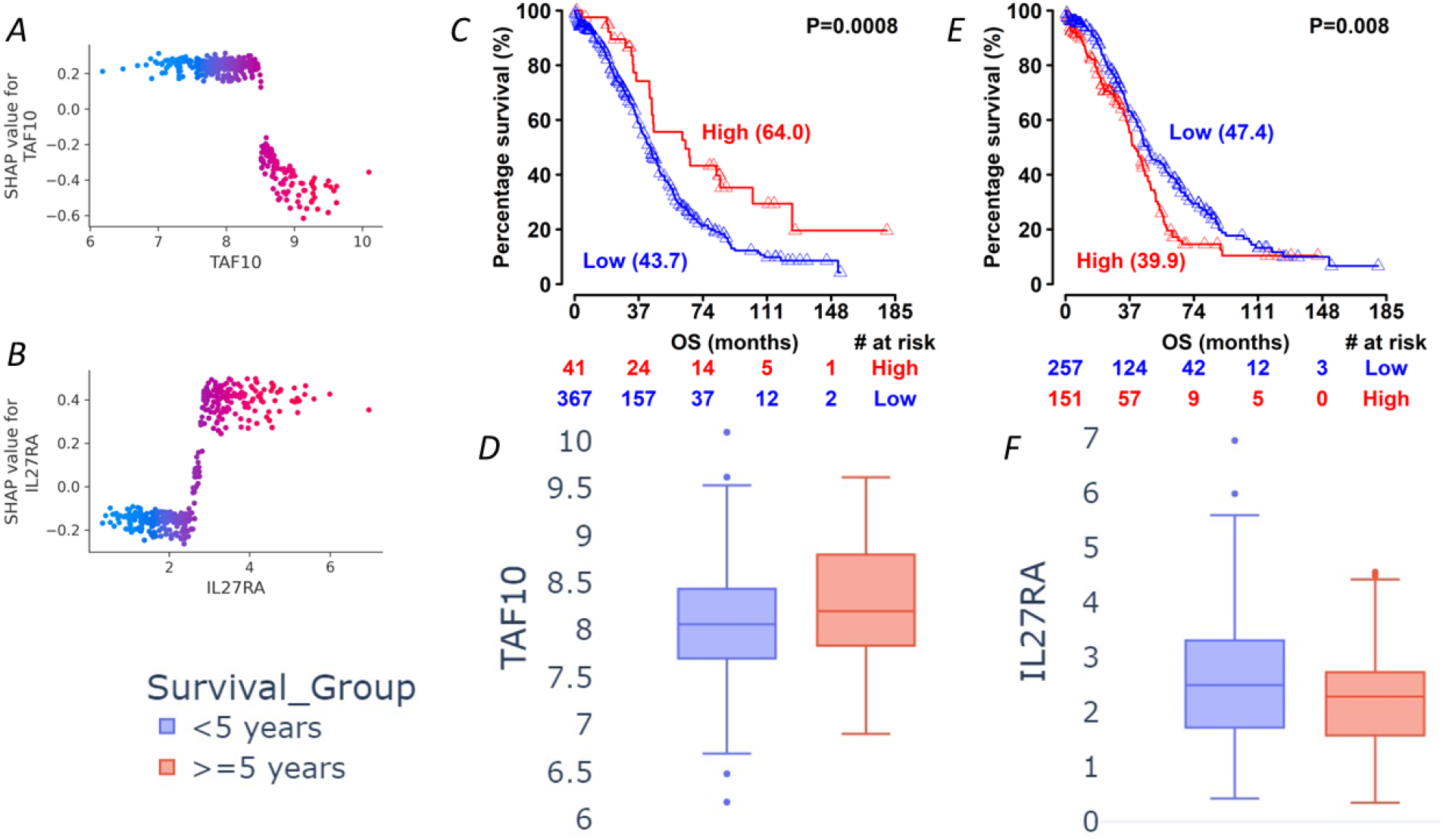
(A) Local biological explanations of most prominent cancer suppressing biomarker (TAF10) gene produced by XAI. (B) Local biological explanations of most prominent cancer promoting biomarker (IL27RA) gene produced by XAI. (C) Kaplan-Meier analysis of TAF10. (D) Box plot analysis of TAF10. (E) Kaplan-Meier analysis of IL27RA. (F) Box plot analysis of IL27RA. (C) & (E) Expression of TAF10 and IL27RA were independent prognostic markers in a cox regression analysis including age and mRNA expression. We used the log-rank test to find the point (cut-off) with the most significant (lowest p-value) split in high/low groups for each gene. The Kaplan-Meier plots were generated for these cutoffs. The numbers of patients at risk in low/high groups at different time points are presented at the bottom of the graph. The analysis and graphing were done in R (http://www.r-project.org/).

